# Efficient Privacy-Preserving String Search and an Application in Genomics

**DOI:** 10.1101/018267

**Authors:** Kana Shimizu, Koji Nuida, Gunnar Rätsch

**Affiliations:** Biotechnology Research Institute for Drug Discovery, National Institute of Advanced Industrial Science and Technology, 2-4-7 Aomi Koto-ku, Tokyo 135–0064, Japan; Information Technology Research Institute, National Institute of Advanced Industrial Science and Technology, 2-4-7 Aomi Koto-ku, Tokyo 135-0064, Japan; Japan Science and Technology Agency (JST) PRESTO Researcher, Tokyo, Japan; Computational Biology, Memorial Sloan Kettering Cancer Center, 1275 York, New York, NY, USA

## Abstract

**Motivation:** Personal genomes carry inherent privacy risks and protecting privacy poses major social and technological challenges. We consider the case where a user searches for genetic information (e.g., an allele) on a server that stores a large genomic database and aims to receive allele-associated information. The user would like to keep the query and result private and the server the database.

**Approach:** We propose a novel approach that combines efficient string data structures such as the *Burrows-Wheeler transform* with cryptographic techniques based on additive homomorphic encryption. We assume that the sequence data is searchable in efficient iterative query operations over a large indexed dictionary, for instance, from large genome collections and employing the (positional) Burrows-Wheeler transform. We use a technique called *oblivious transfer* that is based on *additive homomorphic encryption* to conceal the sequence query and the genomic region of interest in positional queries.

**Results:** We designed and implemented an efficient algorithm for searching sequences of SNPs in large genome databases. During search, the user can only identify the longest match while the server does not learn which sequence of SNPs the user queried. In an experiment based on 2,184 aligned haploid genomes from the 1,000 Genomes Project, our algorithm was able to perform typical queries within ≈4.6 seconds and ≈10.8 seconds for client and server side, respectively, on laptop computers. The presented algorithm is at least one order of magnitude faster than an exhaustive baseline algorithm.

**Availability:** https://github.com/iskana/PBWT-sec and https://github.com/ratschlab/PBWT-sec.

## 1 Introduction

String search is a fundamental task in the field of genome informatics, for which a large variety of techniques have been developed (see, for instance, [2, 17, 20]). Traditionally, those techniques have been optimized for accuracy and computational efficiency, however a recent boom of personal genome sequencing and analyses has spotlighted a new criteria, namely, privacy protection. As reported in many studies, a genome is considered to be one of the most critical pieces of information for an individual’s privacy. In fact, it is largely different from any other personal information because it works as an identifier of an individual while it possesses the information that has strong correlation with the phenotype of the individual [27, 11]. Therefore, in principle, privacy protection is an inevitable problem when handling personal genomes. As a practice, the most popular approach is protecting genomes physically; genomic sequences have been kept at few collaborator sites, and only a limited number of researchers are allowed to access them. This conservative approach severely limits the great potential of existing genomic resources. In order to mitigate the stagnation caused by privacy issues, it appears crucial to develop practical methods that enable searching and mining genomic databases in a privacy-preserving manner.

So far, several groups have tackled related problems. [16] developed secure multi-party computation protocols for computing edit distance. [5] proposed a protocol to search DNA string against a DNA profile represented by finite automata. [6] proposed a protocol to detect a match between two short DNA sequences for the purpose of genetic test. [4] also aimed for genetic test to develop a method for computing set intersection cardinality. [12] proposed a protocol for searching predefined keywords from databases. [24] proposed a substring search protocol for public databases while keeping user’s query private. [3] developed a system by using several cryptographic techniques to find a subset of short reads which includes a fixed-length query string at specific position. [14] proposed an algorithm for finding relatives by secure identity-by-descent matches.

We propose a general approach which utilizes an efficient iteratively queriable data structure together with cryptographic techniques. Among many variations of such data structures, the Burrows-Wheeler Transform (BWT [18, 19, 21]) and related techniques such as the positional BWT (PBWT;[9]) have dramatically improved the speed of genomic database analyses. Those data structures commonly have an indexed dictionary called a rank dictionary. By referring to the rank dictionary in iterative operations, one can efficiently search the database. For the case of BWT, a match between query and database is reported as a left-open, right-closed interval (*f, g*], and the interval is computed by the look-up of the rank dictionary. In our approach, we access the rank dictionary in privacy-preserving manner by using *additive homomorphic encryption* and *oblivious transfer* (OT).

Cryptographic approaches often require significant computational resources. The goal of this work is to illustrate that privacy-preserving queries are within reach when using current cryptographic techniques and standard computing hardware. We demonstrate that a typical query would only take about 4.6 seconds on the user side using a single thread and ≈10.8 seconds on the server having four cores, while preserving privacy of the query string and the database.

The rest of the paper is organized as follows. In Approach, we describe the main ideas of our approach without going into technical details. In Methods, the detailed algorithm of recursive oblivious transfer is given followed by the description of a practical algorithm, named *PBWT-sec*, for privacy-preserving search in large-scale genotype databases. We also describe complexity and security properties of the proposed algorithm. We provide the more intricate details of a more efficient version of the algorithm in Supplementary Sections A-B. In Experiments, we evaluate the performance of *PBWT-sec* on datasets created from data of the 1,000 Genomes Project [30] and compare it to an alternative method for fixed-length *k*-mer search. Finally, we conclude our study in Section 5.

## 2 Approach

### 2.1 Problem Setup

We consider the setting in which a user would like to search a genomic sequence in a database with the aim to either determin whether this sequence exists in the queried database and/or to obtain additional information associated with the genomic sequence. An example is the use in a so-called *genomic beacon* (for instance, those created within the *Beacon Project* of the Global Alliance for Genome&Health (GA4GH).) Another application is the search of a specific combination of variants, for instance, in the BRCA1 or BRCA2 genes, with the aim to determine whether that combination of variants is known or predicted to be deleterious (see, for instance, GA4GH’s *BRCA Challenge*). For privacy reasons, the user would like to conceal the queried sequence, which would be particularly relevant for the second example. For both examples it would be important that the server’s database is protected.

### 2.2 Information Flow of Searches on Recursive Search Data Structures

Let us describe the information flow between a user and a server for such problems. In this work, we perform searches on the (positional) Burrows-Wheeler transform of a genomic database of length *N*. (P)BWT stores string information very efficiently and still allows computations (this is a property of Succinct Data Structures, see [15]).

To search for a query string *q* over the alphabet Σ, one iteratively operates on intervals that can later be used to identify the matching genomic regions based on the (P)BWT. A substring match is represented by an interval (*f, g*]. The number of matches is given by the length of the interval *g* − *f*. It is known that the (*k* + 1)-th interval (*f_k_*_+1_*,g_k_*_+1_] corresponding to a (*k* + 1)-mer match can be updated from the *k*-th interval (*fk,gk*] and the (*k* + 1)-th letter of the query *q*.

We will provide more details on how to update *f* and *g* in Section 3.3. To understand the key ideas, it is sufficient to understand that the updates can be written in the form of

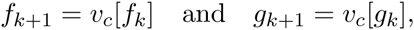

where *c* = *q*[*k* + 1] and *v_c_* ∈ ℕ*^N^* is a large, static lookup table. Hence, the iterative algorithm of updating (*fk,gk*] by using the query *q*, can be written as a recursive algorithm:

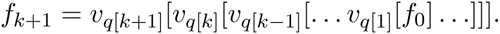

This can be done analogously for *g_k_*_+1_. In this work we will refer to data structures that can be queried in the recursive way described above as *recursive search data structures*. Figure 1 illustrates the information flow of a search on the recursive search data structure.

**Figure 1:**
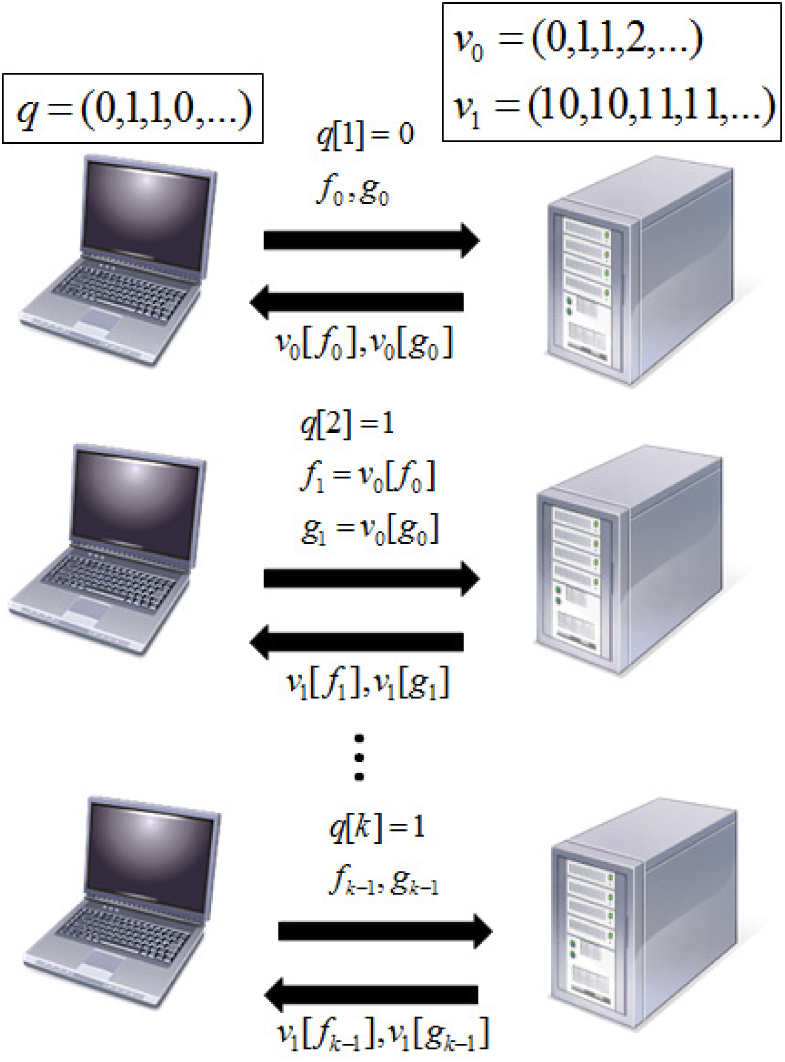
Information flow of a search on a recursive search data structure such as (P)BWT. For *i*-th iteration, the user sends *q*[*i*], *f_i_*_–1_, and *g_i_*_–1_, the server returns *v_q_*_[_*_i_*_]_[*f_i_*–1] and *v_q_*_[_*_i_*_]_[*g_i_*–1], and the user updates *f_i_* = *v_q_*[*i*][*f_i_*_–1_] and *g_i_* = *v_q_*_[_*_i_*_]_[*g_i_*_–1_].

### 2.3 Oblivious Transfer for Privacy-Preserving Search

In a search on the recursive search data structures, the user needs to conceal not only a query string *q* but also *f* and *g* because *f_i_* is *f_i_*_−1_-th element of *v_q_*_[_*_i_*_]_, and *q*[*i*] is inferred from those two values. Analogously, *q*[*i*] is also inferred from *g_i_* and *g_i_*_−1_. The server needs to minimize output because the user reconstructs a part of *v* from the server’s output. In this study, we achieve such security requirements by a cryptographic technique called *oblivious transfer*.

**Oblivious Transfer:** *Oblivious transfer* (OT) is a cryptographic technique for two parties: the user and the server, and enables the user to specify 0 ≤ *t<N* and obtain only *t*-th element of the server’s vector *v* without leaking any information about *t* to the server [26]. Figure 2 illustrates an outline of the oblivious transfer. Among several efficient algorithms [22, 23, 32], we used those which are based on additive homomorphic encryption. The detailed algorithm will be given in Section 3.2.

**Figure 2:**
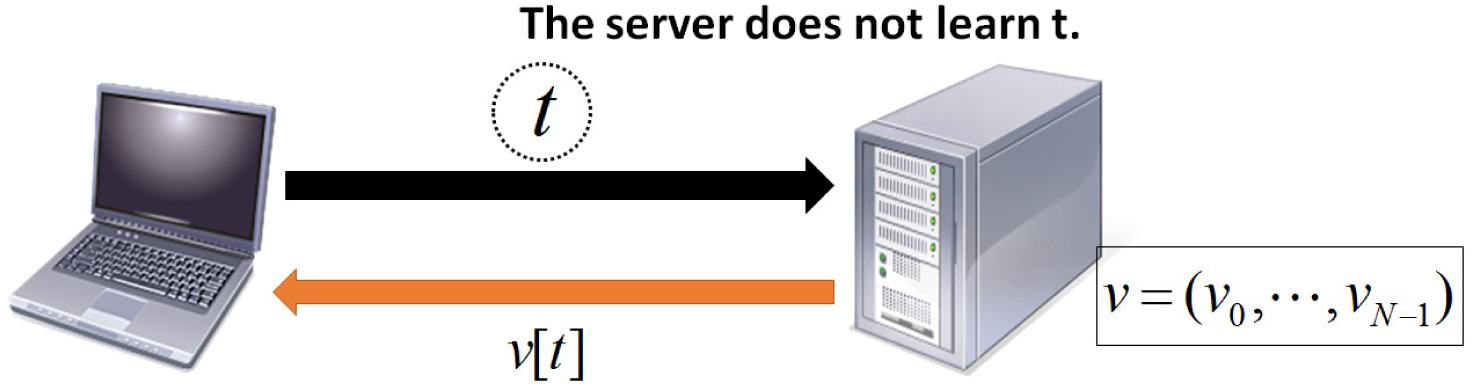
Outline of Oblivious Transfer (OT). The user (laptop computer) obtains *t*-th element of the server’s vector *v* without leaking *t* to the server, and none of the other elements of the vector *v*.

**Concealing the Query:** The user’s query consists of (*f_i_*, *g_i_*], and *q*[*i*+1] for *i*-th iteration. A key idea of our approach is to look-up elements of *v_c_* by OT and obtain the next interval (*f_i_*_+1_ = *v_c_*[*f_i_*], *g_i_*_+1_ = *v_c_*[*g_i_*]] without revealing (*f_i_*, *g_i_*] to the server. In our approach, we also use a masking technique such that the user tries *v_c_*[*f_i_*] for all *c* ∈ Σ, and the server only returns *v_c_*[*f_i_*] where *c* = *q*[*i* + 1] without knowing the value of *q*[*i* + 1]. Technical details will be given in Section 3.2.

**Concealing the Database:** While this approach protects a user’s privacy, the server leaks information of *v_c_* which may be sufficient to reconstruct parts of the genotypes in the database. In order to rigorously protect the server’s privacy, we propose a technique that allows for recursive oblivious transfer where the user does not learn intermediate results but only if a unique match was found. It is based on a bit-rotation technique which enables the server to return 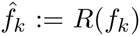 and 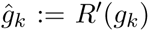 which are random values to the user. Only the server can recover *f_k_* and *g_k_ in encrypted form* (i.e. the server does not see *f_k_* and *g_k_* when recovering them), and thus the user can recursively access *v_c_*[*f_k_*] and *v_c_*[*g_k_*] correctly. The details of this approach are given in Section 3.2.

In this work, we designed an algorithm based on these techniques that can be used for for privacy-preserving search in large genotype databases.

Note that there are still privacy risks for the server, though returning only a unique match minimizes the information leakage from the server. For example, assume there is a database storing a genomic study of drug addicts that implements the *PBWT-sec*, and a person (Bob) participated in the study. If someone obtains a DNA sample from Bob and queries the databases, he/she will reveal that Bob is a drug addict. As described in the above case, there is always a limitation for protecting the server’s privacy as long as the server returns the search results, and there is associated information such as phenotypes [29]. We emphasize that this issue is common for any database search application and is not specific to our proposed method.

## 3 Methods

### 3.1 Additively homomorphic encryption

Our main cryptographic tool in this paper is an additive-homomorphic public-key encryption scheme (KeyGen; Enc; Dec), which enables us to perform additive operations on *encrypted* values. Here, the algorithm KeyGen generates a public key pk and a secret key sk; Enc(*m*) denotes a ciphertext obtained by encrypting message *m* under the given pk; and Dec(*c*) denotes the decryption result of ciphertext *c* under the given sk. The scheme also has the following additive-homomorphic properties:

- Given two ciphertexts Enc(*m*_1_) and Enc(*m*_2_) of integer messages *m*_1_ and *m*_2_, Enc(*m*_1_+*m*_2_) can be computed without knowing *m*_1_, *m*_2_ and the secret key (denoted by Enc(*m*_1_) ⊕ Enc(*m*_2_)).
- Given a ciphertext Enc(*m*) of a message *m* and an integer *e*, Enc(*e* · *m*) can be computed without knowing *m* and the secret key (denoted by *e* ⊗ Enc(*m*)). In particular, Enc(−*m*) can be computed in this manner.

This scheme should have semantic security; that is, a cipher text leaks no information about the original message [13]. For example, we can use either the Paillier cryptosystem [25] or the “lifted”

#### Algorithm 1 Recursive oblivious transfer

1. function PrepQuery(*t*, *N*)
2. *q* = (*q*_0_ = 0;:::; *q_t_* = 1;:::; *q_N_*_–1_ = 0)
3. 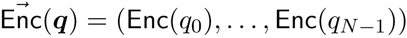
4. return 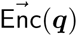
5. end function
6. function 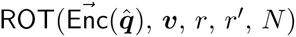
7. 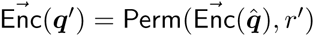
8. 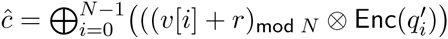
9. return *ĉ*
10. end function
11. *v* is a server’s private vector of length *N*.
12. *x*_1_ is a user’s private value.
13. *x*_ℓ_ is the value of user’s interest.
14. ℓ is known to both user and server.
15. User’s initialization: *t* ← *x*_1_
16. Server’s initialization: *r*′ ← 0
17. Common initialization: *i* ← 1
18. while *i* < ℓ do
19. The user computes: 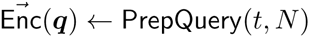
20. if *i* == (ℓ − 1) then
21. Server sets: *r* = 0
22. else
23. Server generates random value *r*
24. end if
25. Server computes: 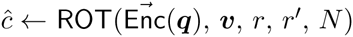
26. Server sets: *r*′ ← *r*
27. Server sends *ĉ* to user
28. User computes: *t* ← Dec(*ĉ*)
29. end while
30. User obtains *x*_ℓ_ = *t*.

version of the ElGamal cryptosystem [10]; now the second operation ⊗ can be realized by repeating the first operation ⊕.

Figure 3 illustrates an outline of performing an additive operation on a user’s value *m*_1_ and a server’s value *m*_2_ by the additively homomorphic encryption. In the first step, the user generates two keys: a secret key and a public key, and the user sends the public key to the server. In the second step, the user encrypts *m*_1_ by the public key and sends a ciphertext Enc(*m*_1_) to the server. In the third step, the server encrypts *m*_2_ by the public key and computes *c* = Enc(*m*_1_ + *m*_2_). The server sends a ciphertext *c* to the user. In the fourth step, the user obtains *m*_1_ + *m*_2_ by decrypting *c*.

**Figure 3:**
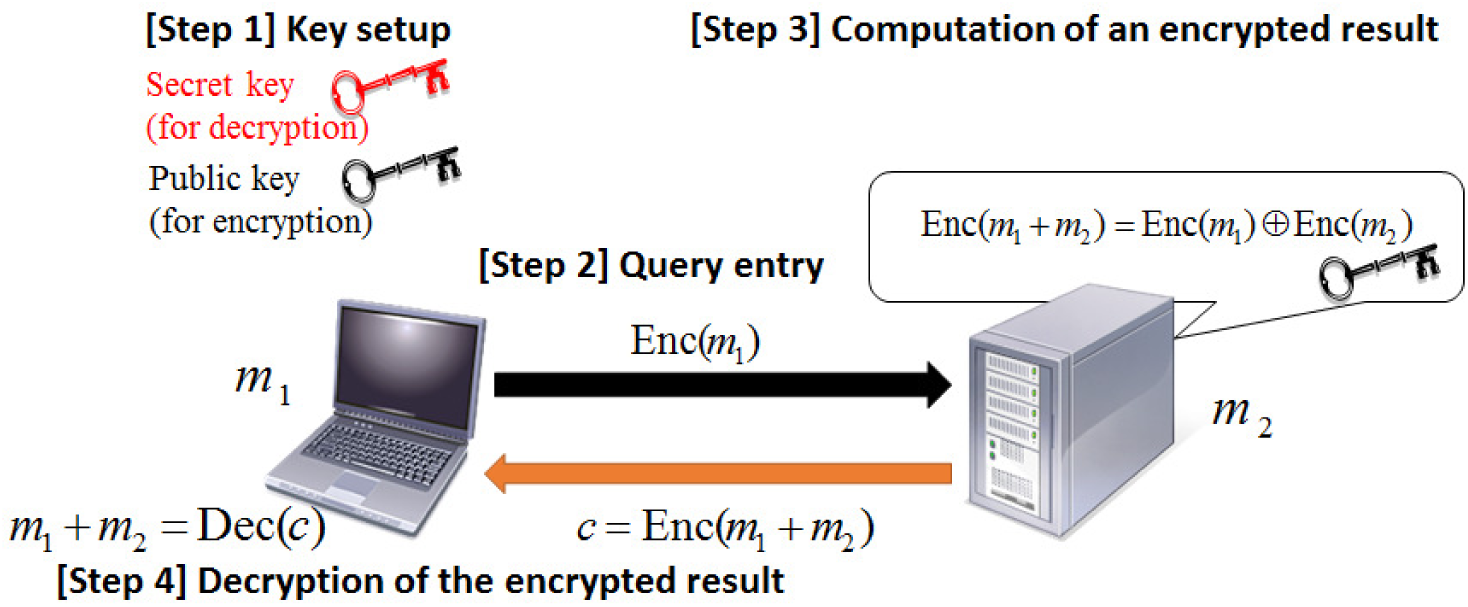
Computation of *m*_1_ + *m*_2_ on the server in encrypted form by additively homomorphic encryption.

It goes beyond the scope of this paper to review the details of these cryptographic techniques and the reader is referred to a book [31] on homomorphic encryption. A typical addition operation in the ElGamal cryptosystem takes about 2·10^−7^ seconds on a single CPU based on AIST’s ElGamal library [1].

### 3.2 Recursive Oblivious Transfer by Random Rotations

To protect the privacy of the database, we propose a technique for recursively querying a data structure without obtaining information about intermediate results. Let us define the recursive oblivious transfer problem as follows:

#### Model 1

*A user has a private value* 0 ≤ *x*1 *<N and a server has a private vector v of length N. Let us denote x_k_*_+1_ = *v*[*x_k_*] *and the user is allowed to access the server ℓ* − 1 *times. After the calculation, the user learns only x_ℓ_ and the server learns nothing about x*_1_*,...,x_ℓ_.*

Here we explain our idea by extending a simple linear communication size OT where the user aims to know the *t*-th element of the server’s vector *v*.

Figure 4 illustrates the oblivious transfer algorithm based on additive homomorphic encryption. In the initialization step, the user generates a public key and a secret key and sends the public key to the server. The user creates a bit vector:

**Figure 4:**
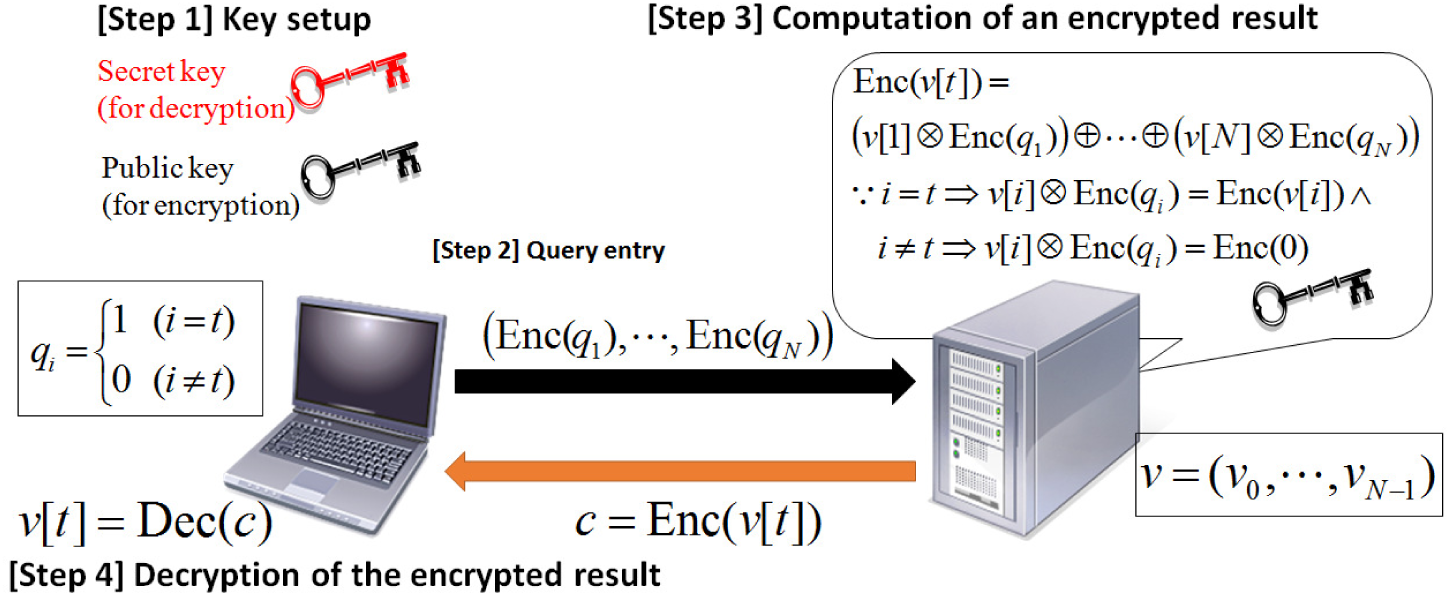
Outline of Oblivious Transfer (OT) based on additive homomorphic encryption. See Sections 2.3 and 3.2 for more details.

**Figure 5:**
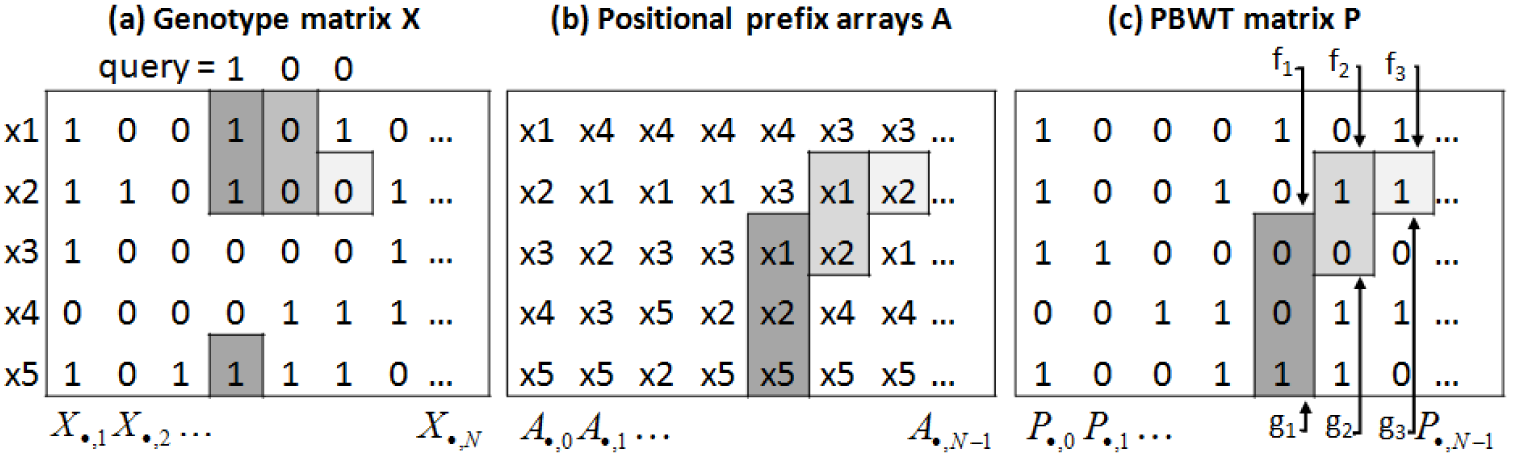
Outline of the search strategy with PBWT. A set of genotype sequences *X* = {*x*_1_*,...,x*_5_} illustrated in (a) is sorted by the algorithm described in [9] to obtain the positional prefix arrays *A* illustrated in (b). Each element *P_i,j_* of PBWT matrix illustrated in (c) is (*j* + 1)-th letter of sequence *A_i,j_*. By computing rank operations with regard to *k*-th query letter on *P*·,_*k*−1_, one can update an interval corresponding to *k*-mer match between the query and *X*. In this figure, the search starts from fourth allele. The first interval (*f*_1_*,g*_1_] is initialized by rank operations on *P*.,_3_ with regard to first query letter ‘1’. The second interval (*f*_2_*,g*_2_] is obtained by rank operations on *P*.,_4_ with regard to the second query letter ‘0’ and (*f*_1_*,g*_1_]. Similarly, the third interval (*f*_3_*,g*_3_] is obtained by rank operations on *P*.,_5_ with regard to the third query letter ‘0’ and (*f*_2_*,g*_2_]. See Sections 2.2 and 3.3 for more details.

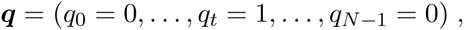

and sends the following encrypted vector to the server.

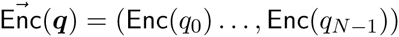

The server computes

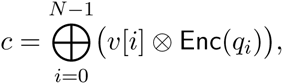

and sends *c* to the user.

The user computes Dec(*c*) and obtains *v*[*t*] by using the secret key because *i* = *t* ⇒ *v*[*t*] ⊗ Enc(*q_t_*) = Enc(*v*[*i*]) and *i* ≠ *t* ⇒ *v*[*t*] ⊗ Enc(*q_t_*) = Enc(0).

Now we consider the case that the server does not leak *v*[*t*], but allows the user to access *v*[*v*[*t*]]. Our idea is that the server generates a random value *r* ∈ {0, 1*,...,N* − 1} and returns the cipher text

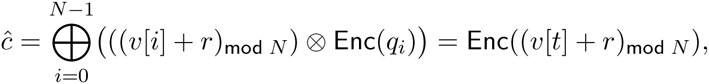

where (*a* + *b*)_mod_ *N* denotes addition in a number field modulo *N*. The user decrypts *ĉ* to know a randomized result (*v*[*t*] + *r*)_mod_ *N*, and performs the next query:

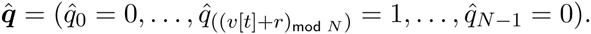

Note that 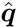 is the *r*-rotated permutation of the ‘true’ query:

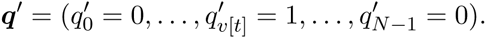

Therefore, denote Perm(*q, r*) as the permutation of a vector *q* such that *i*-th element moves to ((*i* − *r*)_mod_ *N*))-th position, the server can correctly recover ‘true’ query *q′* in its encrypted form by the following permutation: 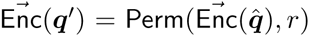. In this way, the server correctly computes an encrypted *v*[*t*]-th element by

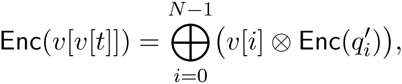

without learning any information about the user’s query.

By recursively applying these calculations, the user can obtain *x_k_*_+1_ according to Model 1. The complete algorithm implementing this idea is given in Algorithm 1. It uses a function ROT for rotating the server’s results to conceal intermediate query results in order to protect the database.

### 3.3 *PBWT-sec*: Privacy-preserving search on genotype databases

In this section, we introduce a practical genotype database search based on recursive oblivious transfer and PBWT. We only introduce the algorithm to search for the longest match starting from *t*-th column, however, variations are possible and would allow for a variety of different search types (see also[9]).

To formulate the problem, let us consider a set *X* of *M* haplotype sequences *x_i_*, *i* = 1*,...,M* over *N* genomic positions indexed by *k* = 1*,...,N*, and a query *q* which is a user’s haplotype sequence over the same *N* genomic positions. We denote *k*-th allele of a sequence *x_i_* by *x_i_*[*k*].

Given two indices *k*_1_ and *k*_2_, we say that there is a match between *q* and *x_i_* from *k*_1_ to *k*_2_, if *q*[*k*_1_] *...q*[*k*_2_ − 1] = *x_i_*[*k*_1_] *...x_i_*[*k*_2_ − 1]. We say that the match is set-longest at *k*_1_ if there is no match between *q* and any sequence *x_j_* (possibly with *j* = *i*) from *k*_1_ to *k*_2_ + 1.

The goal is to find a set-longest match at a given position *t* between *q* and *X* in a privacy-preserving manner. Here, we consider the case that the user’s private information is the query string and the position *t* is not the user’s private information. We later introduce the case that the both the query string and *t* are user’s private information. The formal description of the model is described as follows:

#### Model 2

*The user is a private haplotype sequence holder, and the server is a holder of a set of private haplotype sequences. The user learns nothing but a set-longest match at a given position t between the query and the database while the server learns nothing about the user’s query. t is not a user’s private information and the server knows it.*

Let us remember how to search the set-longest match in *non*-privacy-preserving manner. PBWT involves a matrix *P* ∈ ℕ^*M* × *N*^ that stores well-compressible information in an efficiently searchable form. It is created from the genotype matrix *X* by algorithms described in [9] such that *i*-th column is (*i* + 1)-th letters of sequences sorted by *i* reverse prefix (i.e. sorted from *i*-th letter to first letter). In order to compute the match starting from the first allele, *P* has 0-th column *P*·,_0_ = (*x*_1_[1]*,...,x_M_* [1])*^T^*. By using rank dictionary operations on *P* (see below), one can search a match between a query and *X*. When operating on *P* one computes updates of intervals using the following two quantities (see[9]for more details): i) The rank dictionary for sequence *S* for letter *c* ∈ Σ at position *t*:

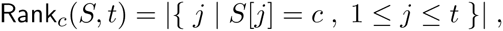

where Σ is the alphabet of *S*. ii) The table CF counting occurrences of letters that are lexicographically smaller than *c* in *S* by

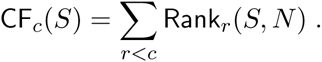

Based on these two quantities, we can compute the updates (*f_k_*_+1_*,g_k_*_+1_] using two simple operations

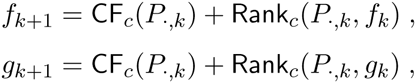

where we denoted the *k*-th column vector by *P*·*_,k_*. Let us define a look-up vector *v_c_* for the column *k* where

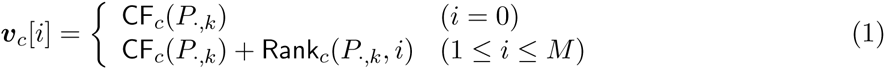

for *c* ∈ Σ. Then, updating an interval is equivalent to two look-ups in the vector *v_c_*:

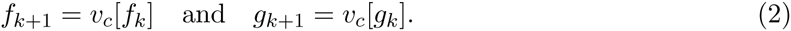

Given a position *t* and a PBWT *P* of the database sequences, the first match is obtained as an interval (*f*_1_ = *v_c_*[0]*,g*_1_ = *v_c_*[*M*]] where *c* = *q*[1] and *v_c_* is a look-up vector for (*t* − 1)-th column of *P* (see the definition of *v_c_* in equation 1). The match is extended by one letter by an update of the interval. The update from the *k*-th interval to (*k* + 1)-th interval is conducted by specifying *c* = *q*[*k* + 1], re-computing *vc* for (*k* + 1)-th column of *P* and referring *v_c_*[*f_k_*] and *v_c_*[*g_k_*] as *f_k_*_+1_ and *g_k_*_+1_ (see (2)). The set-longest-match is found when *f* = *g*.

In order to achieve the security described in the model 2, for each update, the user has to specify *c* without leaking *c* to the server, and obtain only *v_c_*[*f*] and *v_c_*[*g*] without leaking *f* and *g*. To satisfy the second requirement, the user accesses the server’s *v_c_* through the function ROT, which allows the user to obtain a specific element in the specified vector. To achieve the first requirement, the server computes all possible intervals (i.e., computing (*f*, *g*] for the all case of *c* = 0*,...,* |Σ| − 1). This allows the user to obtain the correct interval, however, the sever leaks extra information (i.e., intervals for *c ≠ q*[*k*]). To avoid this, the user sends Enc(*q*[*k*]), and the server adds a conditional randomization factor *r* × (*q*[*k*] − *c*) to *f* and *g* with different random value *r* for all *c* ∈ Σ. Note that this factor becomes equivalent to 0 iff. *q*[*k*] = *c*, and user only obtains the interval for *c* = *q*[*k*].

In order to identify the set-longest match, the user has to know if *f* = *g*. The user cannot compute the identity of *f* and *g* directly from the server’s return, because ROT returns a value which is a random value to the user (but the ‘true’ value is recovered in encrypted form only at the server side). Therefore, the server also sends an encrypted flag *d* which shows whether or not *f* = *g*. Since *f* and *g* are represented as indices of 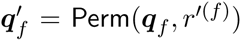 and 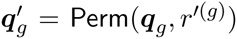 (see the functions PrepQuery and ROT), the server computes *d* by following:

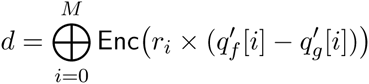

where *r_i_* is a random value. Dec(*d*) is equal to 0 iff. *q_f_* = *q_g_*. See Supplementary Algorithm 7 which defines a function isLongest. In addition to finding a set-longest match at *t*, it is convenient to find a longest substring which matches to at least *ε* sequences. This operation enables to avoid detecting unique haplotype and provides *ε*-anonymity result and is implemented by replacing the function: isLongest by another function: isLongestGT *ε* which computes flags each of which shows if the interval matches to 0*,...,ε* − 1 respectively and returns shuffled flags, and the user knows the result by checking if there is a flag which is equal to zero. See Supplementary Algorithm 7 for more details.

The detailed algorithm of *PBWT-sec* is shown in Algorithm 2.

### 3.4 Concealing the Search Position

By the algorithm introduced above, the match position *t* needs to be provided to the server. Let us consider the case that *t* needs to be concealed (e.g., the used would not like to reveal which gene is analyzed). In practical genotype database search, it is often sufficient for the user to hide *t* in a set of multiple columns. Therefore, here we assume the following security model.

#### Model 3

*The user is a private haplotype sequence holder, and the server is a holder of a set of private haplotype sequences. The user has a vector of D positions T* = (*t*_1_*,...,t_D_*)*. The user learns nothing but a set-longest match at a given position t* ∈{*t*_1_*,...,t_D_*} *between the query and the database while the server learns nothing about the user’s query string. The server knows T but cannot identify which element the user queries.*

Conceptually, the user could query multiple positions at the same time to conceal the search position. In the extreme case the user would query all search positions to avoid leaking any information about *t*. However, every answered query would leak more information from the database and querying would become computationally prohibitive. We therefore propose joint processing using OT that simultaneously uses multiple search positions. Let us define *V_c_* as another look-up vector for a letter *c* as follows:

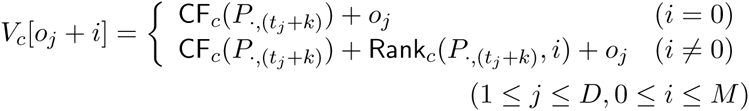

where *o_j_* = (*j* − 1)(*M* + 1) is an offset and *k* is an index which is initialized by −1 and incremented by 1 in each iteration of the recursive search. Note that (*V_c_*[*o_j_*]*,...,V_c_*[*o_j_* + *M*]) corresponds to *v_c_* for *t_j_*-th column. The algorithm for the Model 3 is designed by replacing the lookup tables *v_c_* by *V_c_* (see Step 2a, item 1 in Algorithm 2) and initializing *f* and *g* by *o_x_* and *o_x_* + *M*, respectively, where *t* = *t_x_* (see Step 1 in Algorithm 2). As a result the tables get *D* times larger which has an impact on computing requirements and data transfer size (see Section 3.7). We therefore suggest using this algorithm for small *D*.

### 3.5 Reducing Communication Size

As we will describe in the Complexity analysis in the following section, the *PBWT-sec* algorithm using standard OT requires 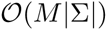 in communication size per iteration in the best case, which makes the core algorithm less practical. We propose to use an algorithm for sublinear-communication OT (SC-OT) proposed in [32]. Using this approach we can reduce the communication size of *PBWT-sec* to 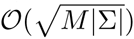 (best case). Here, we only outline the key ideas of SC-OT and its adaptation of *PBWT-sec*. In the SC-OT, the one encodes the position *t* by a two dimensional representation: 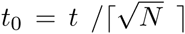, 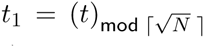, where ⌈·⌉ denotes the ceil of the argument. The user sends Enc(*t*_0_) and 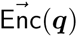 to the server, where

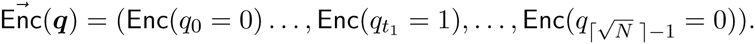

The server obtains random values *r_k_, k* = 0*,...,* 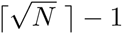, and computes

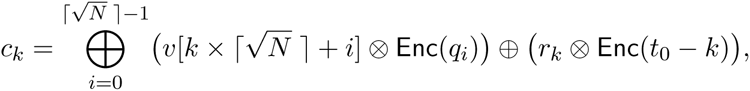

and sends 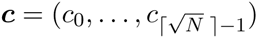 to the user. The user knows the result by the decryption: Dec(*ct*_0_). Note that Enc(*t*_0_ − *k*) = Enc(0) iff. *t*_0_ = *k*, therefore the decryption of *ci* becomes a random value when *i* ≠ *t*_0_.

In order to apply bit-rotation technique naturally to SC-OT, the server needs to return *v*[*t*] in the same two dimensional representation. The key idea here is that the server creates *v*_0_ and *v*_1_ where 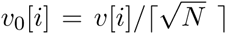 and 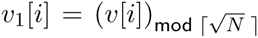, *i* = 0*,...,N* − 1, and searches on both *v*_0_ and *v*_1_. Similar to the linear communication size function ROT, the removable random factors are added to server’s returns. More details on SC-OT is given in Section A. The complete algorithm for privacy-preserving search based on SC-OT is given in Supplementary and Algorithm 4.

### 3.6 An Exhaustive Baseline Algorithm

There are a few related works in regard to finding a DNA substring match [4, 7], however, the goal of *PBWT-sec* is to find the set-longest prefix match from a set of aligned sequences while those works aim to find a fixed-length approximate substring match between two sequences. Therefore, we will compare our algorithm with a baseline algorithm which can find the set-longest prefix match on the basis of exhaustive enumeration of *k*-mers. This baseline algorithm serves as a proxy for the other conceptually similar algorithms.

In order to identify the match, the user queries the server about the presence of a *k*-mer. Here, the server stores all *k*-mers, there are 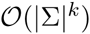 of them, and we use SC-OT. Such a strategy is efficient for short queries as |Σ|*^k^* is not too large. However, the resource requirements will be dominated by queries for large *k* and the algorithm quickly gets intractable.

### 3.7 Complexity

Most of the computing and transfer on server and user side is related to the encryption/decryption and the computational cost of the search is negligible. While PBWT requires essentially 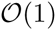 to update the intervals per iteration, *PBWT-sec* needs to conceal the query and requires *M*|Σ| operations on the server, where *M* is the number of sequences in the database and |Σ| is the size of the alphabet. When multiple queries are performed at the same time, i.e., *D* > 1, the effort increases linearly in *D*, i.e., the server sides compute effort is 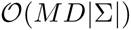 per iteration. When using SC-OT, the communication size and effort for the user is 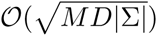 (see Section 3.5 and Supplementary Section A for details).

Table 1 summarizes the time, data transfer overhead and space complexities of the *PBWT-sec*, when the server’s PBWT is *M* × *N* matrix consisting of a set of alphabet letters Σ and the user’s query length is *ℓ* and the number of queries positions is *D* (including *D* − 1 decoy positions; see Section 3.4 for details). For the purpose of comparison, we consider the method outlined in Section 3.6 that achieves the same security and utility as *PBWT-sec*. Since the complexity of the exhaustive approach is exponential to the query length, its performance deteriorates quickly for long matches. On the other hand, the time and data transfer overhead complexity of the *PBWT-sec* are linear and sub-linear to the query length, which enables the user to find a long match efficiently.

**Table 1:** The summary of the time, communication and space complexities of *PBWT-sec* (CP) and an exhaustive method (EX). Both algorithms use SC-OT. *M* is the number of haplotype sequences (server side), *D* is the number of queried positions (including *D* − 1 decoy position to conceal the query position), *ℓ* is the length of query and |Σ| is the alphabet size.

|  | Time | Communication | Space |
| --- | --- | --- | --- |
| CP (user) | $O(\ell\sqrt{MD\vert \Sigma\vert })$ | $O(\ell\sqrt{MD\vert \Sigma\vert })$ | $O(\sqrt{MD\vert \Sigma\vert })$ |
| CP (server) | $O(\ell MD\vert \Sigma\vert )$ | $O(\ell\sqrt{MD\vert \Sigma\vert })$ | $O(MD\vert \Sigma\vert )$ |
| EX (user) | $O(\sqrt{D\vert \Sigma\vert ^\ell})$ | $O(\sqrt{D\vert \Sigma\vert ^\ell})$ | $O(\sqrt{D\vert \Sigma\vert ^\ell})$ |
| EX (server) | $O(D\vert \Sigma\vert ^\ell)$ | $O(\sqrt{D\vert \Sigma\vert ^\ell})$ | $O(D\vert \Sigma\vert ^\ell)$ |

### 3.8 Security Notion

In this paper, we assume the security model called *Semi-honest model* where both parties follow the protocol, but an adversarial one attempts to infer additional information about the other party’s secret input from the legally obtained information. The semantic security of the encryption scheme used in the protocol (see Section 3.1) implies immediately that the server cannot infer any information about the user’s query *q* during the protocol. Also, the user can not infer any information about server’s return except for the result.

Another security model is called *Malicious model* where an adversarial party cheats even in the protocol (e.g., by inputting maliciously chosen invalid values) in order to illegally obtain additional information about the secret. Here we briefly describe one example of an illegal access based on the Malicious model. In our protocol, the user needs to create a bit vector *q* of *N* that includes a bit that is 1 and the rest of the *N* − 1 bits are 0. If the malicious user creates a non-bit vector:

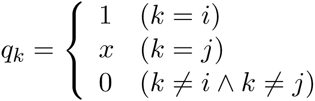

where *x* is a large integer, the server returns *c* = Enc(*v*[*i*] + *x* · *v*[*j*]). When *x* is larger than any element of *v*, the user can infer *v*[*i*] by (Dec(*c*))_mod_*_x_* and *v*[*j*] by Dec(*c*)*/x*. (For example, if *x* = 100 and Dec(*c*) = 821, the user can detect *v*[*i*] = 21 and *v*[*j*] = 8.) Thus the server leaks two elements of *v* by a single query.

In this study, we do not discuss such cases in detail; however, we would like to mention that it is possible to design an algorithm for the Malicious model with a small modification. In order to avoid such attacks, the server needs to verify if the user sends a bit vector which includes only one bit that is 1 and rest of the bits are 0. To achieve this, we suggest using a cryptographic technique called Non-Interactive Zero Knowledge Proofs which enables the user to convince a server that each ciphertext Enc(*m*) corresponds to a value m ∈ {0, 1}, but does not leak any information about which of 0 and 1 is *m*. Among several algorithms, Sakai’s algorithm [28] has such a property. By using the algorithm, the server knows whether or not *q*[*i*] ∈ {0, 1}. To return a correct result only if *q* includes only a single 1, it is sufficient for the server to add *w* = *r* ⊗ Enc(*q*_0_) ⊕ ... ⊕ Enc(*q_N_*_−1_) ⊕ Enc(−1) to the original result, where *r* is a random value. Note that *w* = Enc(0) iff. *q_i_* = 1 and *q_j_* = 0 for 0 ≤ *j* < *N* and *i* ≠ *j*.

## 4 Experiments

In this section, we evaluate the performance of the proposed method on the datasets created from the chromosome 1 data from the 1,000 Genomes Project phase 1 data release which consists of 2,184 haploid genomes [30]. In our experiments and as in [9], we used alleles having SNPs, but we did consider indel variants. We used all 2,184 genomes of original data for all the experiments.

We implemented the proposed algorithm in C++ based on an open source C++ library of *elliptic curve ElGamal encryption* [1]. Our implementation supports communication over the network. We used the standard parameters called *secp192k1* (SECG curve over a 192-bit prime field), according to the recommendation by The Standards for Efficient Cryptography Group. For comparison, we also implemented an exhaustive baseline method (see Section 3.6) that achieves the same security and utility as *PBWT-sec*. In order to perform a fair comparison, both *PBWT-sec* and the exhaustive method used the same SC-OT module where computation of *c_k_* (see Algorithm 1) is simply parallelized by OpenMP.

In the first experiment, the user selected a true start position together with 49 decoys (see Section 3.4 for details), and both *PBWT-sec* and the exhaustive method were run with the same computational setting: the user used a single thread of a laptop computer equipped with an Intel Core(TM) i7 3.00GHz CPU and 16GB memory, and the server used more than eight threads of another laptop equipped with an Intel Core(TM) i7 2.60GHz CPU (four cores with hyper-threading) and 8GB memory. Those two computers communicated over the network.

Figures 6 and 7 show run time and data transfer overhead of *PBWT-sec* and of the exhaustive method. The observed run time and data transfer size of *PBWT-sec* is linear in the query length, while that of the exhaustive approach is exponential. For query lengths larger than 30 bit, the computation of the exhaustive method did not finish within 24 h. These results fit the theoretical complexity described in Section 3.7. We also evaluated performance of the runtime of *PBWT-sec* when the user selected 0, 4, 9, 14, and 49 additional decoy positions. The search with a typical query of length 25 SNP positions and no decoy required no more than 15.5 seconds including communication overhead. (Table 2)

**Table 2:** The run time of a typical query of length 25 SNP positions with *PBWT-sec* on *M* = 2,184 aligned haploid genomes on laptop computers equipped with four cores. The server used all the four cores with hyper-threading while the user used a single thread. All included communication overhead. *D* is the number of positions queried simultaneously to conceal the query position (if required).

|  | User | Server | All |
| --- | --- | --- | --- |
| Run time (sec) with $D = 1$ | 4.55 | 10.8 | 15.5 |
| Run time (sec) with $D = 5$ | 8.77 | 34.0 | 43.0 |
| Run time (sec) with $D = 10$ | 12.5 | 65.3 | 78.0 |
| Run time (sec) with $D = 20$ | 17.3 | 124 | 142 |
| Run time (sec) with $D = 50$ | 27.5 | 311 | 339 |

**Figure 6:**
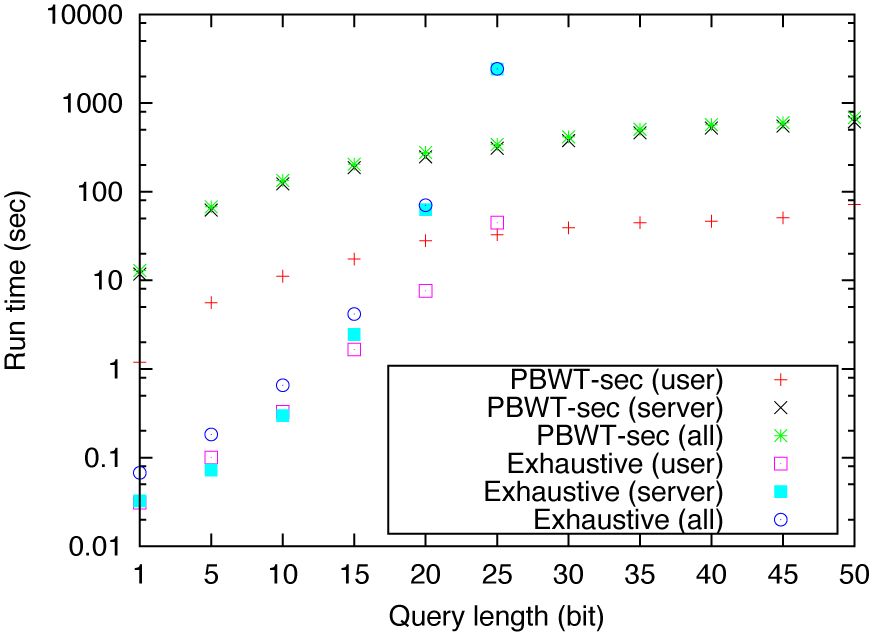
Run time of *PBWT-sec* and the exhaustive method on 2,184 aligned haploid genomes on laptop computers equipped with four cores. The user selected 49 decoy positions for concealing the true query position. The server used all of the four cores with hyper-threading while the user used a single thread. “*PBWT-sec* (all)” and “Exhaustive (all)” include communication overhead.

**Figure 7:**
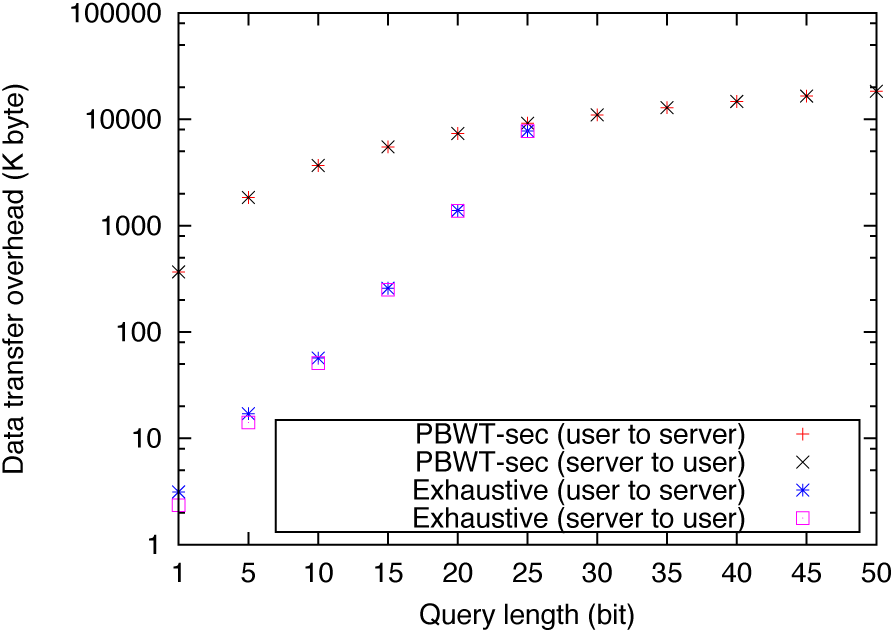
Data transfer overhead of *PBWT-sec* and the exhaustive method on 2,184 aligned haploid genomes on laptop computers. The user selected 49 decoy positions for concealing the true query position.

The user’s run time of *PBWT-sec* is relatively small, making it suitable for a practical case where computation power in a server side is generally stronger than that of user side. Since the memory usage of *PBWT-sec* does not depend on query length, it uses less than 60 MB while that of the exhaustive method exponentially increases according to the query length and required 6 GB when the query length is 25 bit.

Although the exhaustive method is efficient for short queries, we consider that *PBWT-sec* is more practical when taking into account that the bit length of a unique substring for a human genome is greater than 31 bits. Moreover, since there are large linkage blocks, even queries with more than 100 bits would not always lead to unique matches in the 1,000 genomes data. Hence, the exhaustive search strategy would either not always be able to return a unique match or would be very inefficient. The proposed iterative privacy-preserving technique is efficient also for long queries.

In the second experiment, we evaluated the performance of the run time of *PBWT-sec* on a compute node equipped with four CPU sockets (Intel Xeon 2.40GHz CPU; total of 32 cores with hyper-threading). In this experiment, the user also selected 0, 4, 9, 14 and 49 additional decoy positions. For environmental reasons, we did not perform communication over the network and the data was transferred by file I/O which is also included in run time.

Although the current implementation is a prototype and there is room for improvement in terms of parallelization, the server’s run time was at an acceptable level in practical configurations (Table 3). We note, that with improvements in parallelization, the server run time may be reduced to 3–4 seconds.

**Table 3:** The run time of a typical query with *PBWT-sec* on *M* = 2,184 aligned haploid genomes on a compute node with up to 16 cores with hyper-threading and a query length of 25 SNP positions. Wall time includes server (≈90%) and user time (≈10%). *D* is the number of positions queried simultaneously to conceal the query position (if required).

| Parallel Compute Cores | 4 | 8 | 16 |
| --- | --- | --- | --- |
| Run time (sec) with $D = 1$ | 22.6 | 15.5 | 7.9 |
| Run time (sec) with $D = 5$ | 47.3 | 40.0 | 18.4 |
| Run time (sec) with $D = 10$ | 84.5 | 68.4 | 31.6 |
| Run time (sec) with $D = 20$ | 154 | 114 | 56.5 |
| Run time (sec) with $D = 50$ | 386 | 260 | 132.6 |

## 5 Conclusion

In this paper, we have proposed a novel approach for searching genomic sequences in a privacy-preserving manner. Our approach combines an efficient data structure that can be used for recursive search and a novel approach for recursive oblivious transfer. It achieves high utility and has strong security features and requires acceptable compute and communication resources.

The developed novel algorithm can find the longest match between a query and a large set of aligned genomic sequences indexed by PBWT. We implemented our algorithm and tested on the dataset created from the 1,000 Genomes Project data [30]. Compared to an exhaustive baseline approach, our algorithm, named *PBWT-sec*, was orders of magnitude more efficient both in run time and data transfer overhead for practical query sizes. When the prototype program was run on laptop machines, the total run time including communication time over the network was 15.5 sec for searching on 2,184 genomes without concealing the query position. Searches with a concealed query position using a compute node took between 18.6 and 133 seconds depending on the level of privacy.

As the original data structure supports many useful search options such as wild card search and set maximal search, *PBWT-sec* could also support those options by using the same techniques used in the original structures in combination with cryptographic techniques, including OT. Moreover, the approach could be easily applied for BWT and has a potential to be applied for other recursively searchable data structures.

To the best of our knowledge, the proposed algorithm is the first that allows set-maximal search of genomic sequences in a privacy-preserving manner for user and database. We note that the implementation can still be improved and the overall run time can likely be reduced to not more than a few seconds per query. This would make it practical to use our approach in a genomic Beacon (see GA4GH’s Beacon Project) that would allow the privacy-preserving search for combinations of variants. It also appears practical to use our approach to enable search by a user that has access to his/her genomic sequence and would like to query the database, for instance, for information related to disease risk without sharing this information with anybody. Finally, the algorithm can also be used to facilitate sharing of genetic information across institutions and countries in order to identify large enough cohorts with a similar genetic backgrounds. This is in spirit of the mission of the Global Alliance for Genome and Health.

## Acknowledgement

We are thankful to Stephanie L. Hyland for proof-reading the manuscript. We would also like to acknowledge an encouraging discussion with Richard Durbin. We are grateful to the reviewers for their detailed comments that significantly improved the manuscript. We gratefully acknowledge funding from AIST (to K.S.), Memorial Sloan Kettering Cancer Center (to G.R.) and NIH (grant 1R01CA176785-01A1). This study was also supported by the Japan-Finland Cooperative Scientific Research Program of Japan Science and Technology Agency (JST; to K.S.).

## A The sublinear communication size recursive oblivious transfer

In this section, we describe the detailed algorithm of the sublinear communication size recursive oblivious transfer. In Section 3.2, we introduced the bit-rotation technique for the case of the linear communication size oblivious transfer. As mentioned in Section 3.5, the same technique is also applied for the 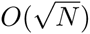-communication size oblivious transfer (SC-OT).

### A.1 The sublinear communication size oblivious transfer

Let us review the SC-OT algorithm. In the SC-OT, the one encodes the position *t* by in a two dimensional representation: 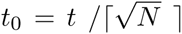, 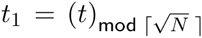, where ⌈·⌉ denotes the ceil of the argument. The user sends Enc(*t*_0_) and 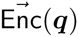 to the server, where

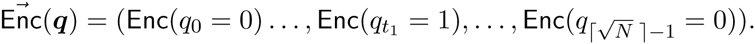

The server obtains random values *r_k_* for *k* = 0*,...,* 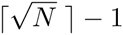, and computes

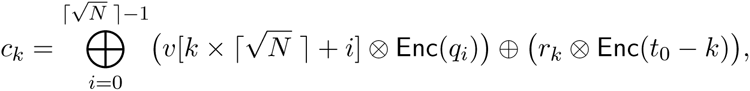

and sends 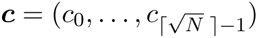 to the user. The user knows the result by the decryption: 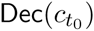. Note that Enc(*t*_0_ − *k*) = Enc(0) iff. *t*_0_ = *k*, therefore the decryption of *ci* becomes a random value when *i* = *t*_0_. See the function SCOT in Algorithm 3 for detailed description.

### A.2 Bit-rotation technique for the sublinear communication size oblivious transfer

In this section, we will describe a new algorithm for the sublinear communication size recursive oblivious transfer (SC-ROT) by using the bit-rotation technique which is introduced in the main text. In order to apply bit-rotation technique naturally to SC-OT, the server needs to return *v*[*t*] in the same two dimensional representation. The key idea here is that the server creates *v*_0_ and *v*_1_ where 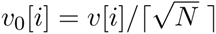 and 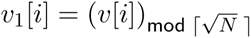, *i* = 0*,..., N* − 1, and searches on both *v*_0_ and *v*_1_. The user obtains next *t*_0_ and *t*_1_ in randomized form by the search on *v*_0_ and *v*_1_ respectively using the same Enc(*t*_0_) and 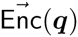. For the search on *v*_0_[*i*], the server generates random value 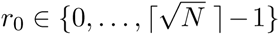 and returns an encrypted value of 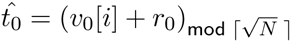. For the search on *v*_1_[*i*], the server generates random value 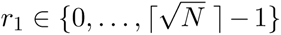 and returns an encrypted value of 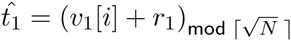. The user decrypts the server’s return and obtains 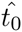 and 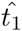 to generate the next query 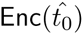 and 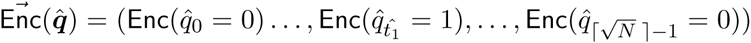. Since the server knows *r*_0_ and *r*_1_, he/she can remove those random factors by 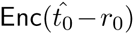 and the circular bit permutation 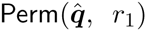 before conducting the next search on *v*_0_ and *v*_1_. To implement such property for the server side, we designed the server’s function SCROT which is described in Algorithm 3. It takes nine arguments: user’s query 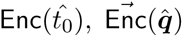, a vector to be searched *v_x_* (*x* ∈{0, 1}), a random value *r* for randomizing the result, upper bound of the true value *L_x_* (*x* ∈{0, 1}), random values 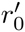 and 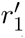 which were used for randomizing ‘true’ values *t*_0_ and *t*_1_ in the previous round (i.e., 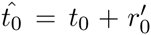 and 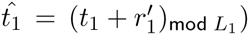) and row length *L*_0_ and column length *L*_1_ of the two dimensional representation (i.e., 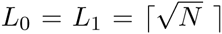 for this case). Figure 8 illustrates the server process for removing random factors previously added to the server’s return. Since 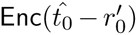 causes the position shift from 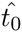 to 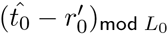 in server’s return *c*, the server also needs another permutation 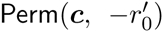 before returning the result. See Algorithm 3 for detailed description. By this function SCROT, the server can add *removable* random factor to the result, and therefore it enables user to search *v* recursively.

**Figure 8:**
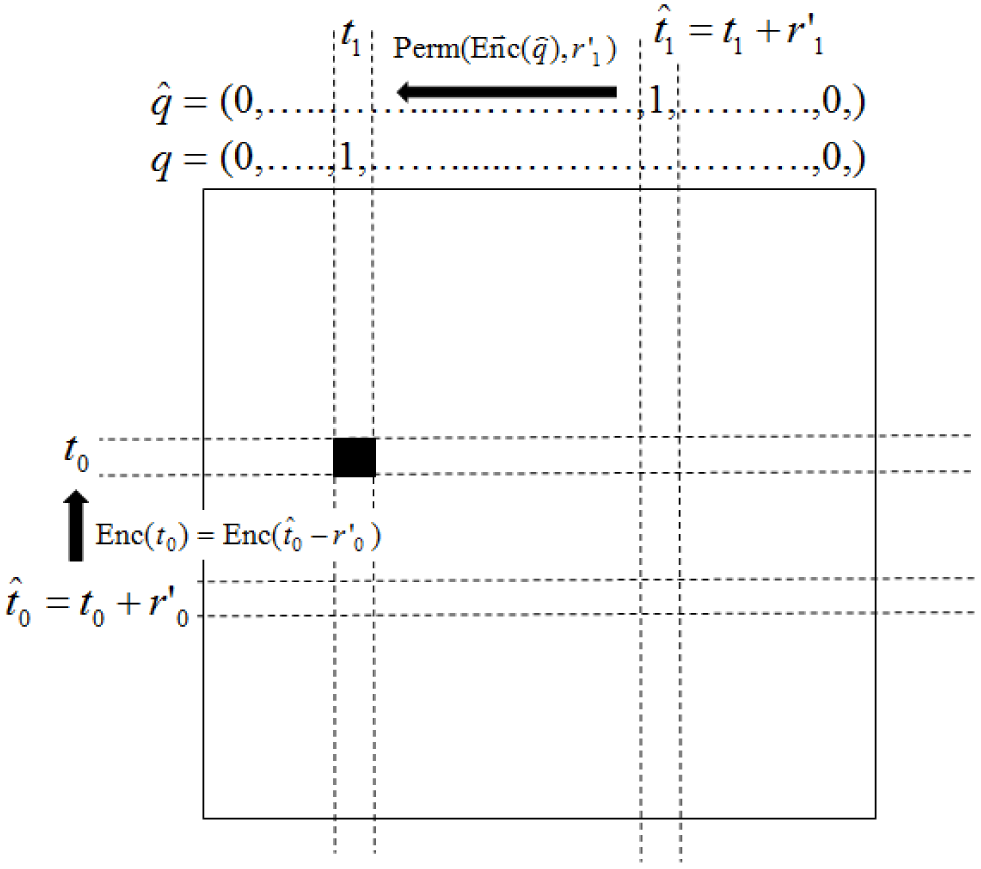
The illustration for the removal of random factors in the server side. *q* and *t*_0_ show the plain text of the user’s ‘true’ query while 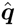 and 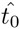 show the plain text of the user’s query. The server recovers correct *t*_1_ by computing 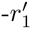 rotated permutation of the server’s query 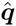. It also recovers correct *t*_0_ by the homomorphic encryption: 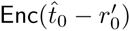.

### A.3 Solving the problem caused by modulo operation of *v*[*i*] + *r*

In the function SCROT, the server generates random value 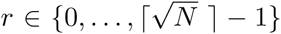 and conducts randomization by:

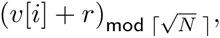

and returns 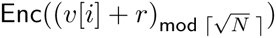 to the user.

Since the modulo operation yields different results for the same *r* according to the two conditions:

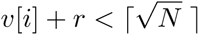

and

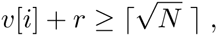

and neither the user nor the server knows which condition is applied (note that the user’s choice *v*[*i*] and server’s random value are their private information), the server needs to return two results assuming both conditions in the next round. For the case of computing Enc(*t*_0_), the sever needs to compute both

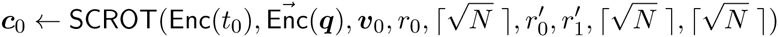

and

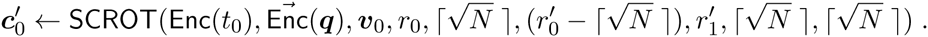

Since only one of 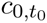 and 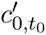 becomes an encryption of a correct result and the other becomes an encryption of a random value, user is able to obtain the next *t*_0_ by checking if 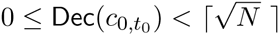 or 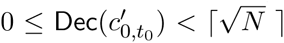 (see the function: ChooseDec in Algorithm 3). In similar way, the user also obtains *t*_1_. Algorithm 4 shows the full description of sublinear communication size recursive oblivious transfer algorithm taking into account of the above problem.

## B The sublinear communication algorithm for *PBWT − sec*

In Section 3.3, the linear size communication algorithm for *PBWT* − *sec* is introduced. Here we introduce the sublinear communication size algorithm by adapting SC-ROT to the search by *PBWT*. The goal is to find a set-longest match at a given position *t* between a query *S* and a set of genotype sequences *X* in a privacy-preserving manner. in this section, we consider that both *t* and *S* are private information and use the following model which is the same model as Model 3 in Section 3.4.

### Model 4

Similar to the linear size communication algorithm for *PBWT* − *sec*, the server creates *v*^(^*^c^*^)^ which is a look-up vector for a letter *c* as follows:

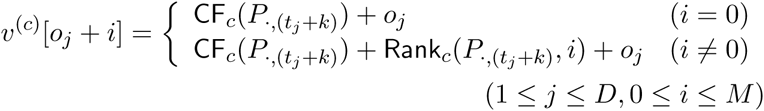

where *o_j_* = (*j* − 1)(*M* + 1) is an offset and *k* is an index which is initialized by −1 and incremented by 1 in each iteration of recursive search. All those letter tables *v^c^* for *c* ∈ Σ are concatenated into one single vector *v* to minimize data transfer overhead. When updating the interval to extend matches by a letter *S*[*i*], the user needs to specify the region of *v*, which corresponds to a letter table *v*^(^*^S^*^[^*^i^*^])^. In our algorithm, we designed row length *L*_0_ and column length *L*_1_ for the two dimensional representation (*L*_0_ and *L*_1_ are not the matrix size of PBWT) such that elements of the same position in the different letter tables should be placed in the same column after concatenating all the tables (i.e., 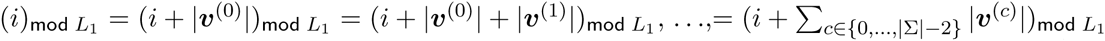) in order that the user can specify the letter table by choosing an offset added to row value (i.e., 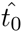) of the query. For this purpose, the server configures 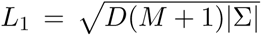, an offset factor 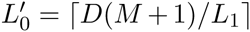, 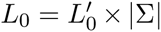, and extend each letter table *v*^(^*^c^*^)^ to the length of 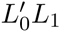 before the concatenation to make *v* (i.e., 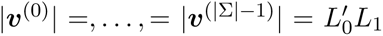). To enable searching *v*^(^*^c^*^)^ by SC-ROT, the server converts all the elements in *v* into the two dimensional representation and stores them in two vectors *v*_0_ and *v*_1_ each of them is of length 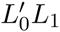. Figure 9 is a graphical view of the rearrangement of *v*_0_ and *v*_1_.

**Figure 9:**
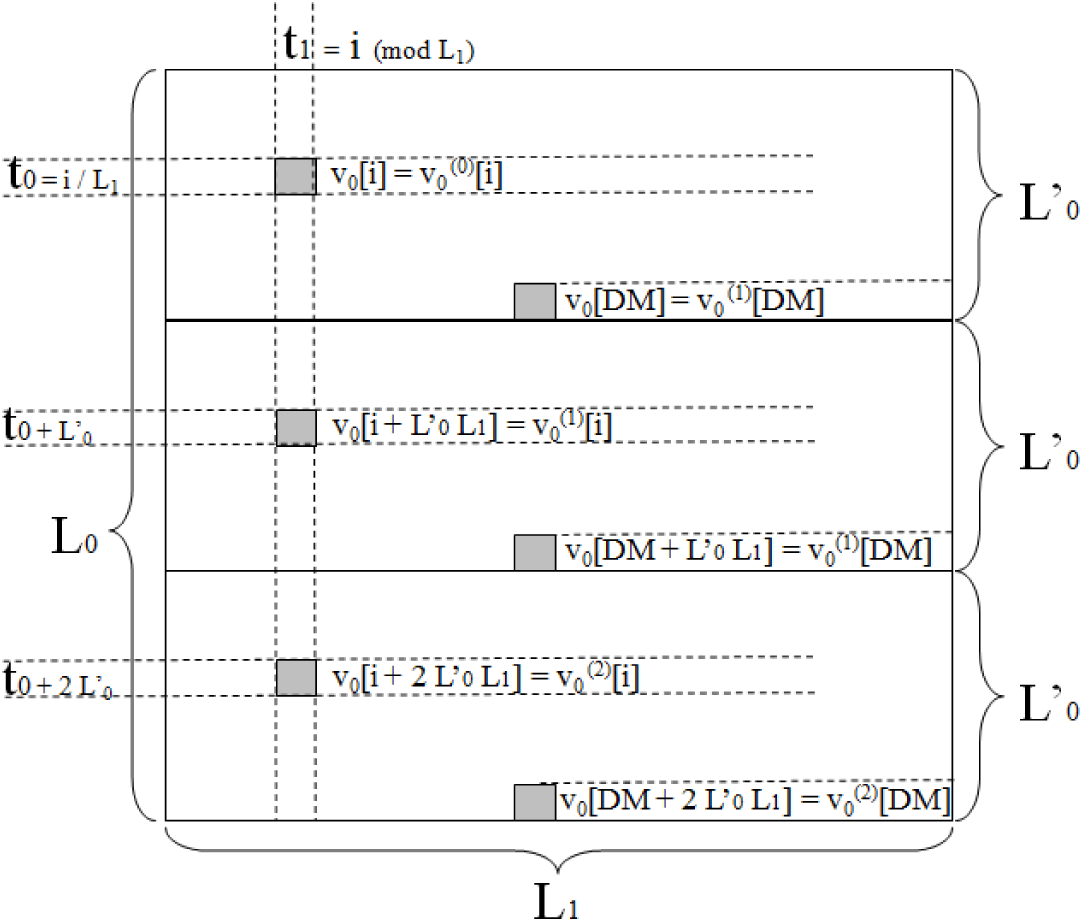
The arrangement of elements of *v*_0_ when Σ = {0, 1, 2}. The length of 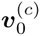 for *c* ∈ Σ is designed such that 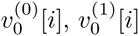 and 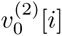 are aligned in the same column after the concatenation. The elements of *v*_1_ is also arranged in the same manner.

Now the user is able to search *v*^(^*^c^*^)^ recursively in an oblivious manner by using SC-ROT. In PBWT, the match is reported as an interval (*f, g*] and the number of matches is equivalent to *g* − *f*. Since the user wants to start the search from *t_x_*-th column on PBWT, user initialized *f* and *g* by *f* = *o_x_, g* = *o_x_* + *M* where *o_j_* = (*j* − 1)(*M* + 1) and computes two dimensional representation of them: *f*_0_ = *f/L*_1_, 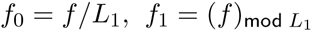, *g*_0_ = *g/L*_1_, 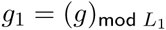. Then the user recursively searches *v*^(^*^c^*^)^ for updating *f* and *g* until he/she finds the match. For the *i*-th round of the recursive search, meaning that the user updates the interval for finding matches ending with *S*[*i*], he/she adds an offset 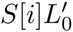 to *f*_0_ and *g*_0_ in order to specify *S*[*i*]. For each round, the server also computes an encrypted flag whose plain text is equal to 0 iff. *f* = *g*. Since there is also a similar problem caused by modulo operation discussed in the section A.3, the server computes the encrypted flag for the case of *v*[*t*] + *r* < *L*_0_ and *v*[*t*] + *r* ≥ *L*_0_. The detailed description of this part is described in the function isSCLongest in Algorithm 5 and item 3-(b) of Algorithm 6. Finally, the user learns the set-longest match at *t* by Dec(*d*). In order to hide the length of the set-longest match to the server, the user keep sending decoy queries until it reaches to *ℓ*-th round. Algorithm 5 and Algorithm 6 show a detailed algorithm of *PBWT-sec*.

### Algorithm 2 The detailed description of *PBWT-sec* finding a set-longest match at position *t*

- Public input: Problem size *M*&*N*; alphabet Σ = {0, 1, .., |Σ| − 1}, start position *t* ∈ {1,...,*N*}
- Private input of user: A query sequence *S* of length *ℓ*
- Private input of server: PBWT matrix *P* ∈ ℕ^*M* × *N*^

0. (*Key setup of cryptosystem*) User generates key pair (pk, sk) by key generation algorithm KeyGen for additive-homomorphic cryptosystem and sends public key pk to server.
1. (*User initialization*) Set initial interval (*f* = 0, *g* = *M*].
2. (*Recursive search*) Initializes query and position index: *i* ← 1; *k* ← *t* − 1 **while** (*i* ≤ *ℓ*) **do**
  a. (*Query entry*) The user performs the following steps:

- Prepares next query:

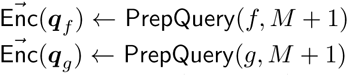
- Sends Enc(*S*[*i*]), 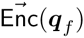, 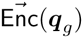 to the server.
  b. (*Search*) The server performs the following steps:

- Compute look-up tables for all *c* ∈ Σ:

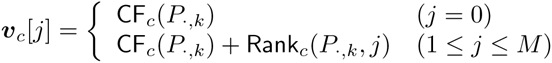
- Obtain random values *r*^(^*^f^*^)^, *r*^(^*^g^*^)^
- Set *r′*^(^*^f^*^)^ = *r*^′(g)^ = 0 iff. *i* == 0
- Compute next possible intervals for all *c* ∈ Σ:

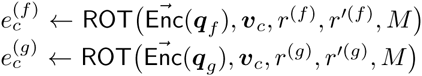
- Randomize return values except for user’s target interval by computing the following for all *c* ∈ Σ Generate temporary random values *r*_0_, *r*_1_

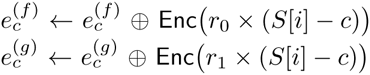
- Compute an encrypted flag showing if match is longest

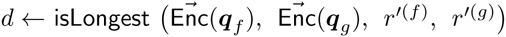
- Store random values *r*′^(^*^f^*^)^ ← *r*^(^*^f^*^)^, *r*′^(^*^f^*^)^ ← *r*^(^*^g^*^)^
- Send *d*, *e*^(^*^f^*^)^, *e*^(^*^g^*^)^ to the user
  c. (*Decryption of encrypted flag and randomized interval*) The user performs the following steps: **if** (Dec(*d*) == 0) Sends decoy queries to server until *i* == *ℓ* Reports result *S*[1,...,*i* − 2] **else** Computes 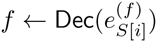, 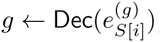 **end if**

*i* ← *i* + 1, *k* ← *k* + 1

**end while**

### Algorithm 3 Building blocks for sublinear communication size recursive oblivious transfer and *PBWT* − *sec*

1. function SCPrepQuery(*t*_0_, *t*_1_, *L*_1_)
2. 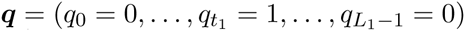
3. 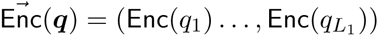
4. return Enc(*t*0), 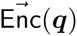
5. end function
6. function SCOT(Enc(*t*0), 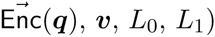
7. for *k* = 0 to *L*_0_ − 1 do
8. Generate random value *r_k_*
9. *x* = *k* × *L*_1_
10. 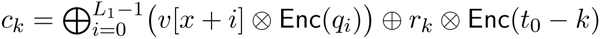
11. end for
12. return 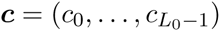
13. end function
14. function 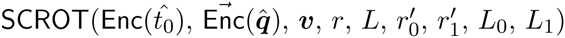
15. 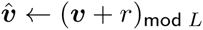
16. 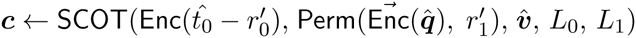
17. 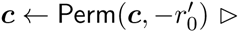 recovering the original position
18. return 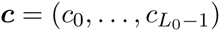
19. end function
20. function ChooseDec(*c*_0_, *c*_1_, *L*)
21. for *x* = 0 to 1 do
22. *m* ← Dec(*c_x_*)
23. if (0 ≤ *m* < *L*)
24. return *m*
25. end if
26. end for
27. end function

### Algorithm 4 The detailed protocol of the sublinear communication size recursive oblivious transfer

- Public input: the database size *N*, query length *ℓ*
- Private input of a user: a start position *t* ∈ 0,..., *N* − 1
- Private input of a server: a vector *v* of length *N*

0. (*Key setup of cryptosystem*) The user generates a key pair (pk, sk) by the key generation algorithm KeyGen for the additive-homomorphic cryptosystem and sends public key pk to the server (the user and the server share public key pk and only the user knows secret key sk).
1. (*Server initialization*) The server computes 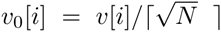, 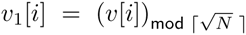 for *i* = 0,...,*N* − 1.
2. (*User initialization*) The user computes 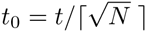, 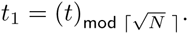
3. (*Recursive search*) Initializes the index by *i* ← 1

**while** (*i* ≤ *ℓ*) **do**

a. (*Query entry*) The user performs the following steps:

- Prepare query **if** (*i* ≠ 1) 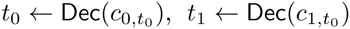 **end if** Enc(*t*_0_), 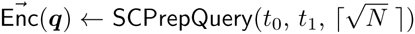
- Sending Enc(*t*_0_), 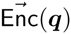 to the server.
b. (*Searching*) The server performs the following steps:

**if** (*i* ≠ *ℓ*)
Generating random values 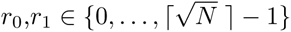
**else**
*r*_1_ = 0, *r*_1_ = 0
**end if**
⊲ ROT removes 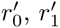 from a query and add *r*_0_ or *r*_1_ to each result.
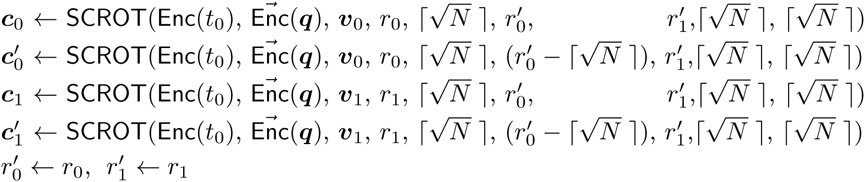

- Sending *c*_0_, 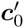, *c*_1_, 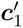 to the user.

*i* ← *i* + 1

**end while**

4. (*Decryption of the result*) The user performs the following steps to obtain result *x*.

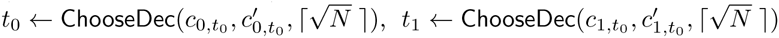

### Algorithm 5 Building blocks for sublinear communication size *PBWT* − *sec*

1. function isSCLongest(Enc(*f*_0_), Enc(*g*_0_), 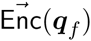, 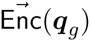)
2. for *i* = 0 to *L*_1_ − 1 do
3. Generating random value *r*
4. 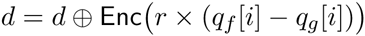
5. end for
6. Generating random value *r*
7. 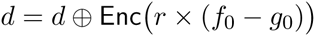
8. return *d*
9. end function
10. function isSCLongestGT*ε*(Enc(*f*_0_), Enc(*g*_0_), 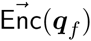, 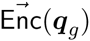, *ε*)
11. for *k* = 0 to *ε* − 1 do ⊲ For the case that *q_g_*[*i*] = 1 move to 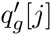 when *i* > *j*
12. 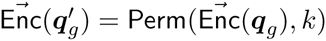 ⊲ 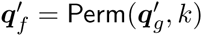 iff. (*g* – *f*) = *k*
13. for *i* = *L*_1_ − *k* to *L*_1_ − 1 do
14. Enc(*q*′*_g_*[*i*]) = Enc(0) ⊲ Avoid a wrong match
15. end for
16. for *i* = 0 to L_1_ − 1 do
17. Generating random value *r*
18. 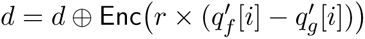
19. end for
20. Generating random value *r*
21. 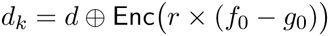
22. end for
23. for *k* = 0 to *ε* − 1 do ⊲ For the case that *q_g_*[*i*] = 1 move to 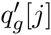 when *i* < *j*
24. 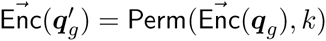 ⊲ *q*′*_f_* = Perm(*q*′*_f_*, *k*) iff. (*g* – *f*) = *k*
25. for *i* = 0 to *k* do
26. Enc(*q*′*_g_*[*i*]) = Enc(0) ⊲ Avoid a wrong match
27. end for
28. for *i* = 0 to *L*_1_ − 1 do
29. Generating random value *r*
30. 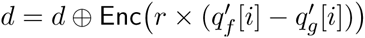
31. end for
32. Generating random value *r*
33. 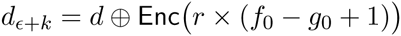
34. end for
35. *d* = (*d*_0_,...,*d*_2_*_ε_*)
36. Shuffling order of elements in *d*
37. return *d*
38. end function

### Algorithm 6 The detailed description of sublinear communication size *PBWT − sec* for finding a set-longest match at position *t*

- Public input: The length of column *M*, a set of alphabet letters Σ = {0, 1, .., |Σ|− 1} and a set of (*D* − 1) decoy positions and true position *T* = (*t*_1_,...,*t_D_*).
- Private input of a user: A starting column *t_x_* ∈ *T*, a query sequence *S* of length *ℓ*
- Private input of a server: PBWT matrix P ∈ ℕ^*M* × *N*^

0. (*Key setup of cryptosystem*) The user generates a key pair (pk, sk) by the key generation algorithm KeyGen for the additive-homomorphic cryptosystem and sends public key pk to the server (while only the user knows secret key sk).
1. (*Server initialization*)

- The server computes 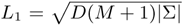, 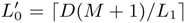, 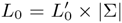 and announces L_0_, L_1_ and 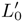 to the user.
2. (*User initialization*)

- The user initialize a half-open interval (*f*, *g*] by *f* = *o_x_*, *g* = *o_x_* + *M* where *o_j_* = (*j* − 1)(*M* + 1).
- The user computes two dimensional representation of (*f*, *g*] by *f*_0_ ← *f*/*L*_1_, 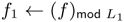, *g*_0_ ← *g*/*L*_1_, 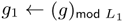
3. (*Recursive search*) Initializes the indices by *i* ← 1 *k* ← − 1 while (*i* ≤ ℓ) do
  a. (*Query entry*) The user performs the following steps:

- Prepare next query: 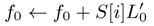, 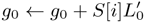 ⊲ Setting offset to search matches ending with *S*[*i*]

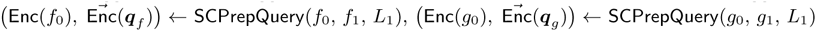
- Sending Enc(*f*_0_), 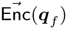, Enc(*g*_0_), 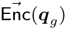, Enc(*S*[*i*]) to the server.
  b. (*Searching*) The server performs the following steps:

- Computes vectors *v*^(^*^c^*^)^ of length *D* × (*M* + 1) for all *c* ∈ Σ:

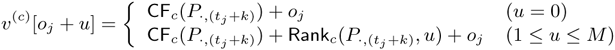

where *o_j_* = (*j* − 1)(*M* + 1) for *j* = 1,...,*D*.
- Creates vectors 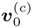, 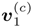 of length 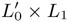 for *c* = 0,..., |Σ| − 1.
- Computes 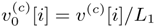, 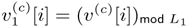 for *i* = 0,..., *D*(*M* + 1) − 1 and *c* = 0,..., |Σ| − 1.
- Creates vectors *v*_0_ and *v*_1_ by concatenating 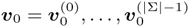 and 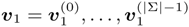.
- Generates random values 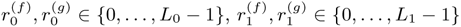
- Computes next intervals and an encrypted flag showing if the match is the longest

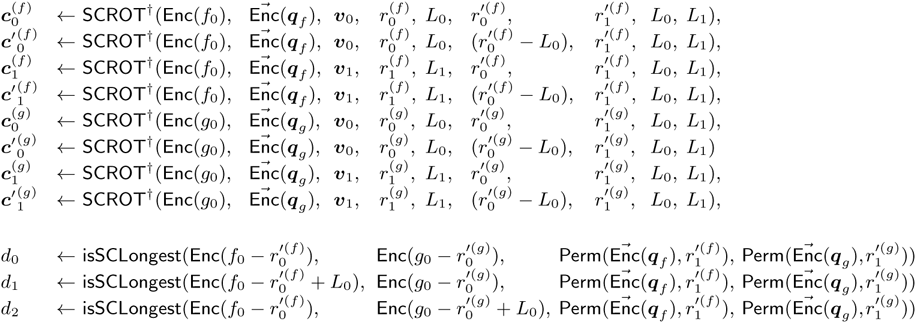
- Storing random values 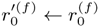, 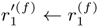, 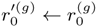, 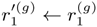
- Sending 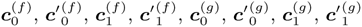, *d* to the user
  c. (*Decryption of the encrypted flag and the randomized interval*) The user performs the following steps: if (Dec(*d*_0_) == 0 || Dec(*d*_1_) == 0 || Dec(*d*_2_) == 0) Reports the result *S*[1,..., *i* − 2] and sending the server decoy queries until i == ℓ else Computes 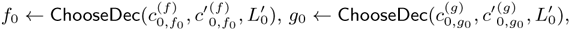 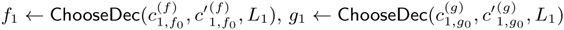 ⊲ for choosing correct results end if

*i* ← *i* + 1 *k* ← *k* + 1

end while

### Algorithm 7 Building blocks for linear communication size *PBWT* − *sec*

1. function 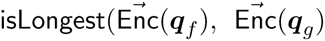, *r*′^(^*^f^*^)^, *r*′^(^*^g^*^)^)
2. 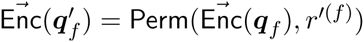
3. 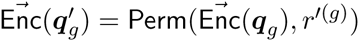
4. for *i* = 0 to *M* do
5. Generating random value *r*
6. 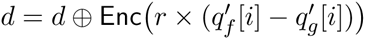
7. end for
8. return d
9. end function
10. function 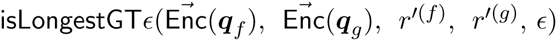
11. for *k* = 0 to *ε* – 1 do
12. 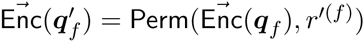
13. 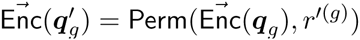
14. 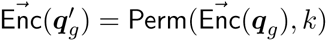 ⊲ *q*′*_f_* = Perm(*q*′*_f_*, *k*) iff. (*g* – *f*) = *k*.
15. for *i* = 0 to *M* do
16. Generating random value *r*
17. 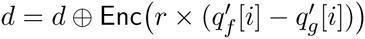
18. end for
19. *d_k_* = *d*
20. end for
21. d = (*d*_0_, …, *d_ε_*)
22. Shuffling order of elements in *d*
23. return *d*
24. end function

